# ChIPSeqFPro, a pipeline for sequential processing of ChIP-Seq fastq to bigwig files

**DOI:** 10.1101/118281

**Authors:** Milos Pjanic

## Abstract

ChIPSeqFPro is a pipeline that automates processing of a collection of ChIPSeq or ATAC-Seq data starting from the gzip compressed fastq files. It performs the quality control using FastQC, mapping to the human genome hg19 or mouse mm10 using BWA mapper, for both single read or paired end sequencing fastq files, followed with sam to bam conversion using samtools view, creates statistics on bam files using samtools flagstat, peak calling with MACS, and finally creates high resolution bigwig files from bam files using a custom script bam2bigwig that invokes bedtools bamtobed and UCSC scripts, bedItemOverlapCount, bedGraphToBigWig and fetchChromSizes.

## Introduction

Processing ChIP-Seq or ATAC-Seq sequencing data (1–3) requires multiple intermediary computational steps. ChIPSeqFPro (ChIP-Seq Full Processing) is a pipeline that will perform sequential processing of ChIPSeq data starting from the compressed fastq files, i.e. fastq.gz files. It performs FastQC (4) quality control, mapping to the human genome hg19 or mouse genome mm10 using BWA mapper (5) for fastq files that originate from single read or paired end sequencing experiments, sam to bam conversion using samtools view (6), obtaining statistics on bam files using samtools flagstat (6), peak calling with MACS (7), and finally creates bigwig files (8) from bam files using a custom bam2BigWig tool. Bam2BigWig will convert bam files to bigwig files for viewing in the UCSC Genome (9) or WashU (10) browsers. Bam2bigwig depends on bedtools bamtobed (aka bamToBed) (11), and it will attempt to download and execute three UCSC scripts, bedItemOverlapCount, bedGraphToBigWig and fetchChromSizes (Figure 1). ChIPSeqFPro will process all fastq.gz files that are placed in a working folder and in case of paired end sequencing it will pair the correct R1 and R2 fastq.gz files for subsequent processing.

**Figure 1.**
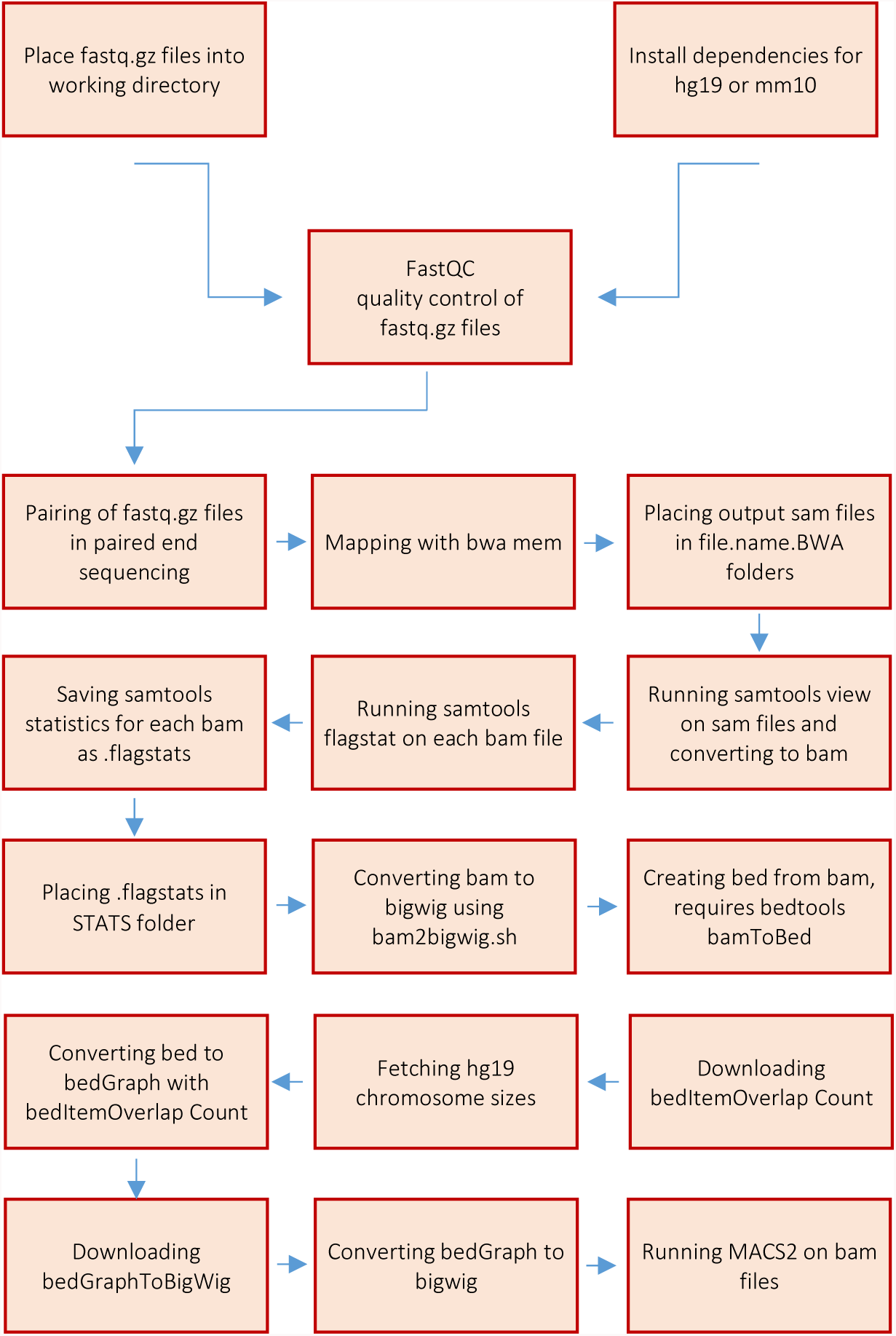
Schematic representation of the ChIPSeqFPro pipeline. ChIPSeqFPro pipeline involves FastQC quality control, mapping to the human genome hg19 or mouse mm10 using BWA, for both single read or paired end sequencing, sam to bam conversion using samtools view, creates statistics on bam files using samtools flagstat, peak calling with MACS, and finally creates bigwig files from bam files using a custom script bam2bigwig that invokes bamToBed and bedItemOverlapCount.

ChIPSeqFPro consists of four bash scripts, ChIPSeqFPro.PE.hg19.sh, ChIPSeqFPro.PE.mm10.sh, ChIPSeqFPro.SR.hg19.sh, ChIPSeqFPro.SR.mm10.sh, that will process single read sequencing files on human genome 19 (ChIPSeqFPro.SR.hg19.sh), single read sequencing files on mouse genome 10 (ChIPSeqFPro.SR.mm10.sh), paired end sequencing files on human genome 19 (ChIPSeqFPro.PE.hg19.sh) and paired end sequencing files on mouse genome 10 (ChIPSeqFPro.PE.mm10.sh).

## Dependencies

Place fastqc.gz file collection for processing in a working folder.

~~~
mkdir work.folder
cp path-to-files/*fastq.gz work.folder
~~~

**FastQC instalation (Linux). Place FastQC folder in a working directory.**

~~~
cd work.folder
wget http://www.bioinformatics.babraham.ac.uk/projects/fastqc/fastqc_v0.11.5.zip
unzip fastqc_v0.11.5.zip
chmod 755 ./FastQC/fastqc
~~~

**Reference genome installation. Download the reference genome, in this example the human genome, hg19.**

~~~
mkdir ~/reference_genomes
cd ~/reference_genomes
mkdir hg19
cd hg19
wget --timestamping
     ’ftp://hgdownload.cse.ucsc.edu/goldenPath/hg19/bigZips/hg19.2bit'
     -O hg19.2bit
wget http://hgdownload.cse.ucsc.edu/admin/exe/linux.x86_64/twoBitToFa
chmod 755 twoBitToFa
./twoBitToFa hg19.2bit hg19.fa
~~~

**Installation for the mouse genome, mm10.**

~~~
mkdir mm10
cd mm10
wget --timestamping
   ’http://hgdownload.cse.ucsc.edu/goldenPath/mm10/bigZips/mm10.2bit'
   -O mm10.2bit
./twoBitToFa mm10.2bit mm10.fa
~~~

**Install BWA. Download to your home (~) folder the latest version of BWA from http://sourceforge.net/proiects/bio-bwa/files/**.

~~~
bunzip2 bwa–0.7.15.tar.bz2
tar xvf bwa–0.7.15.tar
cd bwa-0.7.15
make
~~~

**Edit ~/.bashrc to add BWA to your PATH**.

~~~
nano ~/.bashrc
export PATH=$PATH:~/bwa-0.7.15
source ~/.bashrc
#test the installation
bwa
~~~

**Indexing the reference genome with BWA. Use BWA to index the reference genome.**

~~~
cd ~/reference_genomes
#for human genome
bwa index -a bwtsw hg19.fa
#for mouse genome
bwa index -a bwtsw mm10.fa
~~~

**Installing MACS2 through PyPI system.**

~~~
pip install MACS2
~~~

**Download bam2bigwig for human genome.**

~~~
wget https://raw.githubusercontent.com/milospjanic/bam2bigwig/master/bam2bigwig.sh
chmod 775 bam2bigwig.sh
~~~

**Alternatively, download bam2bigwig for mouse genome.**

~~~
wget
https://raw.githubusercontent.com/milospjanic/bam2bigwig/master/bam2bigwig.mm10.sh
chmod 775 bam2bigwig.mm10.sh
~~~

## Usage

ChIPSeqFPro is composed of four pipelines that will run on either human genome hg19 or mouse genome mm10, using either paired-end (PE) or single-read (SR) sequences, ChIPSeqFPro.PE.hg19.sh, ChIPSeqFPro.PE.mm10.sh, ChIPSeqFPro.SR.hg19.sh, ChIPSeqFPro.SR.mm10.sh. After placing fastq.gz files in the working folder run the script that corresponds to the sequencing type and species, e.g:

~~~
chmod 755 ChIPSeqFPro.PE.hg19.sh
./ChIPSeqFPro.PE.hg19.sh 64
~~~

The first argument indicates number of cores you want to allocate for the analysis, use the number of cores on your machine, e.g. 64. In order for script to properly work, it is necessary to place the FastQC folder into the working folder. This is the only requirement necessary to be placed in the working directory, in addition to the collection of fastq.gz files that are being processed.

In addition, the reference genome has to be places to be in your ~/reference_genomes folder, therefore in case you switch to another user account script may not work because it searches for the home folder of the current user, (~/reference_genomes.

## Repository

The code for all four ChIPSeqFPro scripts could be downloaded from the GitHub repository, https://github.com/milospianic/ChIPSeqFPro. and the code for two bam2bigwig scripts could be downloaded from the GitHub repository, https://github.com/milospianic/bam2bigwig.

## References

1. Pjanic M, Pjanic P, Schmid C, Ambrosini G, Gaussin A, Plasari G, et al. Nuclear factor I revealed as family of promoter binding transcription activators. BMC Genomics. 2011;12:181.

2. Pjanic M, Schmid CD, Gaussin A, Ambrosini G, Adamcik J, Pjanic P, et al. Nuclear Factor I genomic binding associates with chromatin boundaries. BMC Genomics. 2013;14:99.

3. Miller CL, Pjanic M, Wang T, Nguyen T, Cohain A, Lee JD, et al. Integrative functional genomics identifies regulatory mechanisms at coronary artery disease loci. Nat Commun. 2016 Jul 8;7:12092.

4. Babraham Bioinformatics - FastQC A Quality Control tool for High Throughput Sequence Data [Internet]. [cited 2016 Dec 7]. Available from: http://www.bioinformatics.babraham.ac.uk/projects/fastqc/

5. Li H, Durbin R. Fast and accurate short read alignment with Burrows-Wheeler transform. Bioinforma Oxf Engl. 2009 Jul 15;25(14):1754–60.

6. Li H, Handsaker B, Wysoker A, Fennell T, Ruan J, Homer N, et al. The Sequence Alignment/Map format and SAMtools. Bioinforma Oxf Engl. 2009 Aug 15;25(16):2078–9.

7. Zhang Y, Liu T, Meyer CA, Eeckhoute J, Johnson DS, Bernstein BE, et al. Model-based analysis of ChIP-Seq (MACS). Genome Biol. 2008;9(9):R137.

8. Kent WJ, Zweig AS, Barber G, Hinrichs AS, Karolchik D. BigWig and BigBed: enabling browsing of large distributed datasets. Bioinformatics. 2010 Sep 1;26(17):2204–7.

9. Kent WJ, Sugnet CW, Furey TS, Roskin KM, Pringle TH, Zahler AM, et al. The Human Genome Browser at UCSC. Genome Res. 2002 Jun 1;12(6):996–1006.

10. Zhou X, Maricque B, Xie M, Li D, Sundaram V, Martin EA, et al. The Human Epigenome Browser at Washington University. Nat Methods. 2011 Dec;8(12):989–90.

11. Quinlan AR, Hall IM. BEDTools: a flexible suite of utilities for comparing genomic features. Bioinformatics. 2010 Mar 15;26(6):841–2.

